# A deep learning workflow for quantification of Micronuclei in DNA damage studies in cultured cancer cell lines: a proof of principle investigation

**DOI:** 10.1101/2022.09.18.508405

**Authors:** Anand Panchbhai, Munuse Ceyda Ishanzadeh, Smarana Pankanti, Ahmed Sidali, Nadeeen Solaiman, Radhakrishnan Kanagaraj, John J Murphy, Kalpana Surendranath

## Abstract

The cytokinesis block micronucleus assay is widely used for measuring/scoring/counting micronuclei, a marker of genome instability in cultured and primary cells. Though a gold standard method, this is a laborious and time-consuming process with person-to-person variation observed in quantification of micronuclei. We report in this study the utilisation of a new deep learning workflow for detection of micronuclei in DAPI stained nuclear images. The proposed deep learning framework achieved an average precision of >90% in detection of micronuclei. This proof of principle investigation in a DNA damage studies laboratory supports the idea of deploying AI powered tools in a cost-effective manner for repetitive and laborious tasks with relevant computational expertise. These systems will also help improving the quality of data and wellbeing of researchers.

**Simple Summary:** This study aims to test a suitable deep learning method for micronucleus detection in images acquired for cytokinesis block micronucleus assay. This study has reached a mean average precision of >90%.

## 1. Introduction

DNA damage induced genomic instability is a defining characteristic of human cancers and several other diseases and therefore is at the centre of biochemical, molecular, and clinical investigations of human diseases [1]. By products of cellular metabolism, replication errors and external sources such as environmental factors and chemotherapeutic drugs promote damage to the double helix. The array of cellular responses activated post DNA damage and outcomes of cell division provide clues about proper cell function and faithful transmission of the genome to daughter cells. Failure to recognise and remove DNA defects is associated with disease pathogenesis, particularly in oncogenesis. Micronuclei (MN) are extranuclear bodies originating from chromosome fragments or whole chromosomes that lag in anaphase during cell division. A common effect of mitotic dysfunctions that consequently lead to whole chromosome mis-segregation during cell division is MN [2–6]. MN are not included in the daughter nuclei and serve as valuable biomarkers to evaluate genome integrity. However, experiments require a strenuous process of scoring thousands of binucleated cells and manual counting of MN either directly through the microscope or from the images. This method to study genome instability in a reproducible and reliable manner is time and resource demanding.

Cell detection and tracking has been an age-old problem since the inception of microscopes. Visual inspection by eye has been the common method for detection and counting of cells which is often time consuming and prone to human errors. With the advent of technology, computer vision and automation are being introduced in every aspect of research. Extensive research validates the application of Artificial Intelligence (AI) in detection of cells infected in malaria, tuberculosis, and several use cases [7–15]. However, the adoption of these methodologies in the assessment of DNA damage responses have been comparatively limited. Several research labs around the world depend on manual counting for analysis of genome instability data. This lack of adoption of AI tools may be attributed to lack of awareness, unavailability of necessary resources and a scepticism about the abilities of AI’s in general. Answering these factors forms the motivation of this study, where we try to show the efficacy of a deep learning system and elaborate on the cost benefits of adopting such a system in a laboratory setting. Automation and streamlining the studies of genome instability is what the authors believe holds the potential to accelerate the work being done in research labs by many folds.

## 2. Related Work

ML has been used in the domain of medical sciences to cater to several use cases ranging from COVID-19 detection from chest X-rays [11] to diabetic retinopathy [12]. Efficient methods to study DNA damage as well as integration of ML has the potential to yield reliable results and decrease turnaround times in low-resource academic research and hospital settings. Proof of concept genotoxic studies [16–24], have tried to capitalize on several statistical models to detect nuclei; these studies have been of limited use due to lack of differentiation between cell types for detections [19]. Studies have attempted to use off-the-shelf ML algorithms and tools like Cell Profiler to perform segmentation of cell nuclei in microscopic cell images [20]. Caicedo et al. suggested that the algorithms they tested faced difficulties in detecting tiny objects like MN as they tended to be confused with debris and artefacts which comprise a good portion of objects [20]. Bahreyni Toossi et al. [24] attempted to detect increased frequency of micronuclei (abbreviated as MNi in the article) on Giemsa-stained inverter CBMN assay. This four-staged process which exhibited a sensitivity of 82% involved the segmentation of nuclei, detection of the binucleated cell (BN) followed by estimation of the whole cell with an assumption of non-existent cytoplasm and the detection of MNi within each BN. Recently, a convolutional neural network was deployed to detect mononucleated cells, binucleated cells, polynucleated cells, mononucleated cells with MN [17].

Interestingly, it was apparent that deep learning models tended to perform well for detecting and segmenting nuclei of varying sizes and shapes in microscopic images [19]. Several studies [17,24] focused on detecting cells followed by identification of MN. The results of manual scoring showed that such an approach may lead to underestimation of the number of MN [25]. Therefore, the authors of current study use object detection for the scoring of MN instead of a two staged method used in earlier studies. In this study, a deep learning-based object detection model was used to detect MN in microscopic images. Object detection deals with detection of instances of semantic objects in digital images and includes both localization and classification of objects in an image. The novel aspects of this paper also include the release of a large, annotated dataset of CBMN assay images (available upon request to corresponding authors), which the authors believe will help in future studies in this domain and identification of new AI powered methods for the detection and counting of MN from microscopic CBMN assay images.

## 3. Materials and Methods

### 3.1 Preparation of slides and the dataset

#### 3.1.1 Cell lines and culture conditions

Osteosarcoma cell line U-2 OS (American Type Culture Collection, HTB-96™) and U-2 OS T-Rex used in this study were grown as monolayers in T25 and T75 flasks at 37°C with 5% CO2 in a humidified atmosphere. Cells were cultured in Dulbecco’s modified Eagle’s medium supplemented with 10% fetal bovine serum and 1% penicillin-streptomycin (P/S) antibiotic (Tetracycline-free serum). Cells were passaged with trypsin and viability was assessed with 0.4% trypan blue solution and the cell density was determined using Denovix Cell DropTM cell counter. Images utlised in the study were obtained from micronuclei experiments of 2 doctoral researchers in the genome engineering laboratory.

#### 3.1.2 Reagent for DNA damage assay

Aphidicolin, the DNA polymerase inhibitor was purchased from Sigma and dissolved in DMSO at a concentration of 3 mM.

#### 3.1.3 Micronucleus assay

U-2 OS wild type and T-Rex variants were grown on Poly-D-Lysine coated coverslips (Corning, #354086) in 12-well plates and individually treated with three concentrations [0.1 μM, 0.2 μM, 0.4 μM] of aphidicolin for 24 hrs. Cell culture medium was supplemented with 2 μg/ml of cytochalasin B (Sigma-Aldrich, #C6762) 16 hrs prior to fixation to block cells at cytokinesis stage of cell division. Cells were fixed with PTEMF buffer [20 mM PIPES (pH 6.8), 10 mM EGTA, 0.2% triton X-100, 1mM MgCl2 and 4% formaldehyde] for 10 minutes at RT and mounted with prolong gold antifade mounting medium containing DAPI. Slides were dried, and edges of coverslips sealed and stored at 4C until further analysis

#### 3.1.4 Scoring of MN

Slides were analysed with an upright fluorescent microscope (Olympus BX41) equipped with an Elite Micropix camera (Micropix Ltd). Images were acquired at 100x using Cytocam software Version 2.0 (Micropix). Over 1600 DAPI-stained binucleated cells were analysed and distinct MN in the vicinity of these cells were scored manually. Binucleated cells were scored for MN in the presence and absence of aphidicolin, which is extensively used to assess replication stress in cultured cells.

#### 3.1.5 Manual Annotation of MN

A dataset containing 1642 images were manually annotated for the study and will be released publicly upon request. (CBMNA2021). The images were captured at a magnification of 100 x with an overall magnification of 1000x. Bounding boxes were used as a method of data annotation as shown in figure 1. The online tool of Logy.ai was used and a total of three annotators annotated each image. The annotators were requested to enclose the whole micronuclei within the bounding box without including the surrounding objects. There was no restriction imposed on the size or the number of bounding boxes. A detection was considered as a ground truth only if two or more annotators agreed to avoid discrepancies in this methodology. The annotators were doctoral researchers skilled in detecting micronuclei and had at least 2 years of experience.

**Figure: 1:**
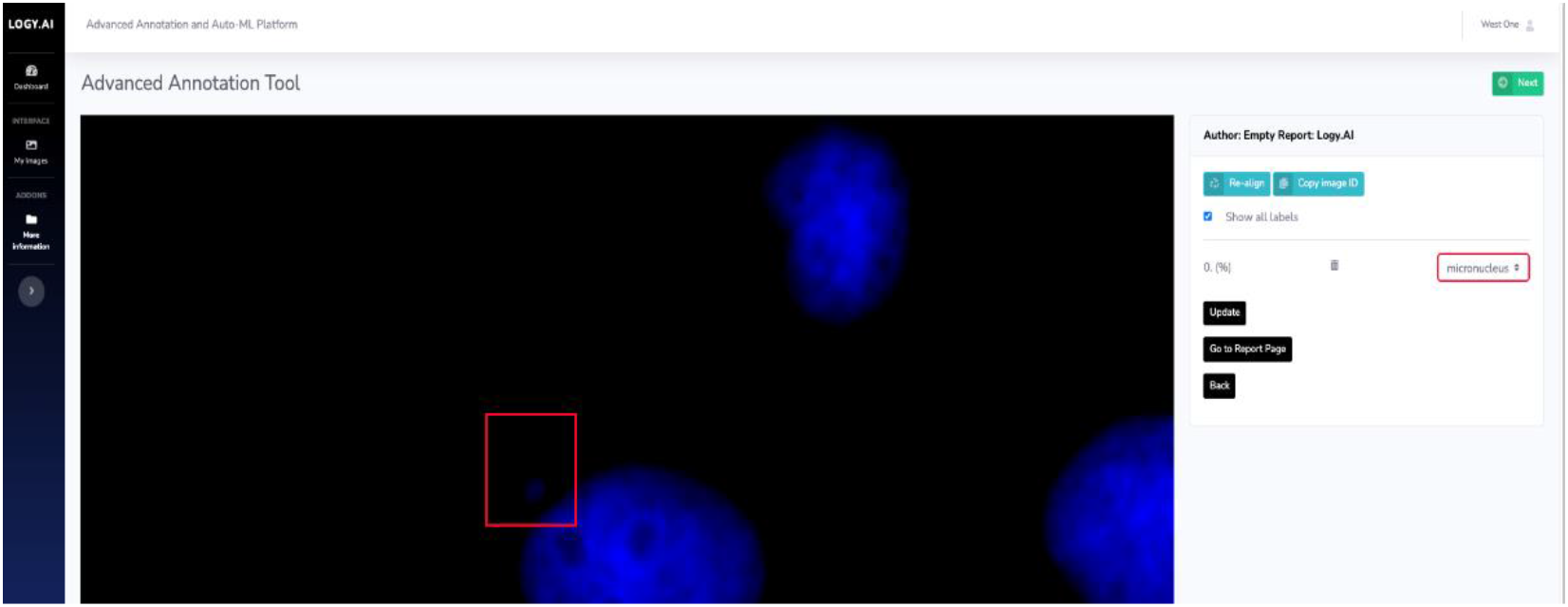
Image of Logy.AI annotation portal used for annotating micronuclei. The annotators were required to draw boxes over the image. Each box was then assigned to a class using the right panel. Only one class was used in this study: micronucleus.

### 3.2 Deep learning workflow

The overall methodology can be broken down into multiple steps, (step 1) these steps include the preparation of the slides to be observed under a microscope and capturing the images of the slides at a certain magnification, (step 2) annotating the captured images, (step 3) training the AI models based on reference annotation on a part of the annotated data (Train and Validation Set) followed by (step 4) inference results from the AI models on a completely new set of microscopic images (Test Set).

Logy.AI Auto ML platform (LaiAMLP v1.0) was used as part of this study. It is a cloud based end-to-end no-code platform for building ML models. LaiAMLP is a scalable web server built on python-3.7 and NGINX which uses TensorFlow v1.15 to train deep learning models based on the input data and corresponding ground-truth. LaiAMLP can be accessed over any standard browser like Chrome, Safari or Edge. The intuitive interface of LaiAMLP is specifically designed to eliminate the need of any technical expertise. Post preparation of data (step 1), it is uploaded to LaiAMLP where it can then be staged for annotations (step 2). LaiAMLP provides a simple drag and drop interface to create boxes on the regions of interest, here MN as shown in figure 1c.

Once the data is annotated it is split into training, validation and testing sets as per the requirements. LaiAMLP then initiates the ML training process (step 3) and continuously provides learning statistics and on-the-go annotations on the test set at a regular number of epochs and produces a readily deployable ML model as a TensorFlow-Serving-Container which is a snapshot of the learned parameters that can then be immediately used for generating (step 4) inferences. These ML models can be easily integrated into the workflow of a data heavy laboratory processing 1000s of images per second.

#### 3.2.1 FasterRCNN

Faster RCNN runs on a unified and more economical framework and shares the results of feature extraction with both the region proposal and classification steps. This decreases the computational need without a trade-off in performance. Additionally, it can be more accurate as it uses the same CNN for both region proposal and object detection, avoiding errors that can be introduced by different architectures. The RPN is responsible for generating a set of potential object regions in the image, called region proposals. These regions are then passed to the Fast RCNN Classifier, which is responsible for classifying each region as containing an object or not. The RPN outputs a set of region proposals. It works by sliding a small CNN window, called an anchor, over the entire feature map, and for each location and generates multiple region proposals, each with a different scale and aspect ratio. Each region proposal is then passed through the Fast RCNN Classifier, which is also implemented as a CNN. The detector takes the region proposal and a feature map extracted from the image, and it applies a set of convolutional and pooling layers to extract features from the region proposal. These features are then passed through a fully connected layer that generates a score for each class, indicating the likelihood that the region proposal contains an object of that class. The reason for selecting Faster RCNN for this study was due to its established superiority in detection of the smaller objects [27]. Micronuclei are often smaller in size hence this architecture was selected as per the results of Huang et al. [27]

##### Data Flow

Faster RCNN (figure 2.0) has two stages: *region proposal network (RPN)* and *Fast RCNN Classifier* [26]. The image is first resized while maintaining the aspect ratio with a minimum dimension of 600 and a maximum dimension of 1024 with all the 3 colour channels. The resized image (A) is then fed into a Resnet-101 [26,28] feature extractor (B) to create a feature map extracted from an intermediate layer of Resnet-101 of size 14×14 (C). A number of feature extractors are available publicly. Resnet 101 [28] was used as this architecture and feature extractor combination is known to work well for small-sized detections [27]. The extracted features boxes are used to predict class-independent box proposals (D). The number of box proposals was set to 100, in order to achieve a balance between speed and accuracy [26,27]. The box proposals are used to crop features from the feature map (E). The cropped feature maps are then fed to the softmax classifier which predicts the class of the box and provides refinements related to position and size of the green boxes in F. Each box proposal is rated between 0-1 for each class which is called the objectness score. The class with the highest value is assigned to that box proposal. Furthermore, a minimum threshold is also used as an eligibility criterion to classify a box proposal, the selection of these thresholds is based on the value that gives the best performance. The green boxes in F depict the boxes which got classified into available classes while the other box proposals were discarded. These classified boxes are then mapped on the image to show the detections (G). As the features are not computed twice for these two stages, Faster RCNN [26] runs faster than its counterparts [27], hence the name.

**Figure 2:**
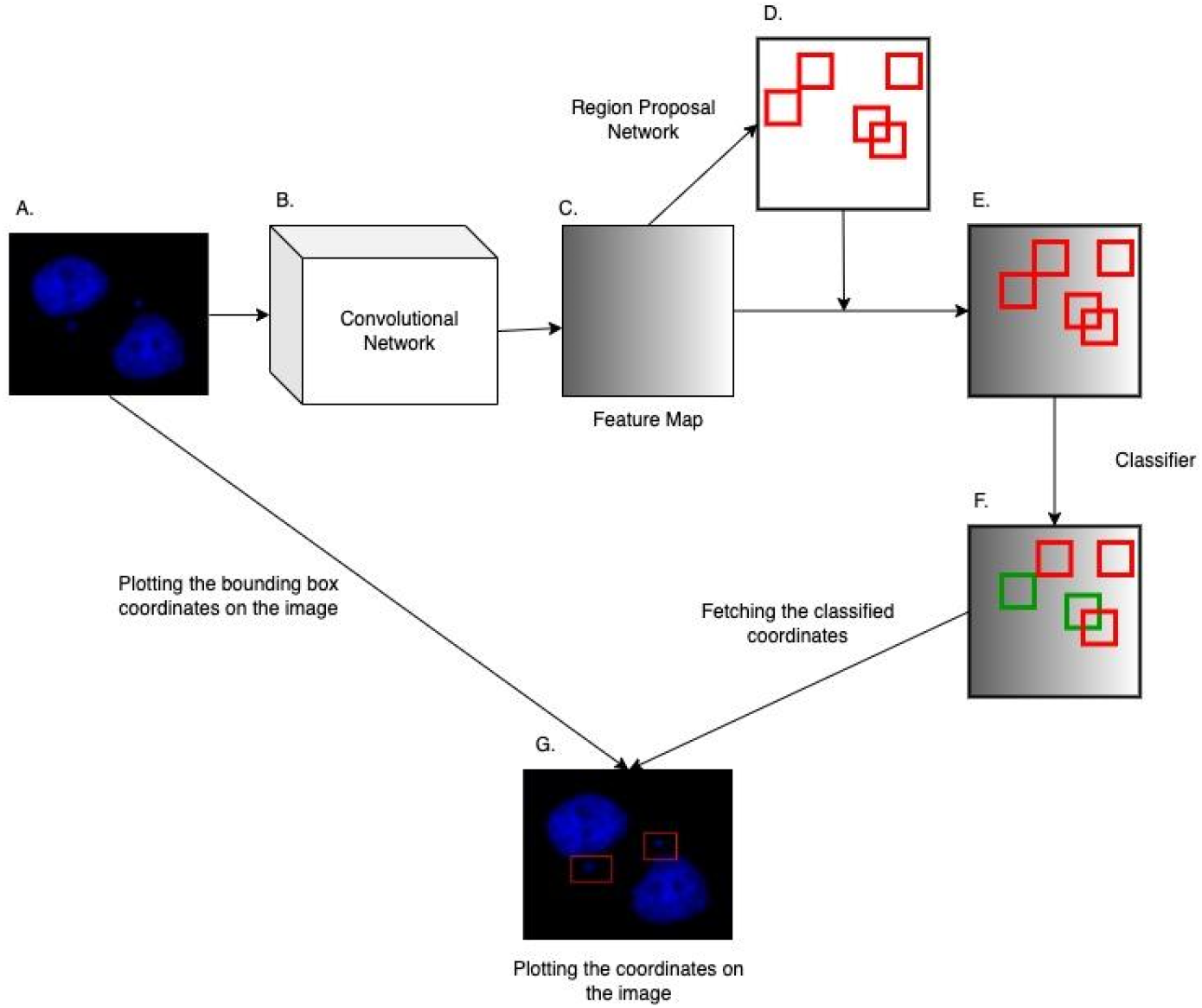
Faster RCNN flow diagram. Faster R-CNN [26] consists of two main components: a Region Proposal Network (RPN) and a Fast RCNN Classifier.

#### 3.2.2 Training Configuration

Data augmentation is used to improve the robustness of the model. The images of the dataset are rotated, flipped and cropped before feeding into the training phases. This allows the model to understand the morphology of the object being detected irrespective of its angle or position. For this study, random horizontal flips were used for data augmentation. It was deemed enough due to the general random distribution of micronuclei in an image. A gradient descent optimizer with momentum was used for the training process for support in noisy data distribution and for overcoming shallow local minima. Learning decay was also used due to its efficiency in process of finding local minima and preventing overshooting that often takes place in the training process. A variable learning rate also assisted us to speed up learning process in the initial phases where a higher learning rate is used, and a lower learning rate is used towards the end to emulate fine adjustments.

### 3.3 Training the AI model

A dataset containing 1642 manually annotated images was used for this study (CBMNA2021). The images were captured at a magnification of 100 x with an overall magnification of 1000x. The training was carried out by LaiAMLP. The model used for this study is a modified version of Faster-RCNN coupled with coco-resnet-101 [26] for feature extraction. The images annotated contained only one label namely micronucleus. The training was performed on a TensorFlow (a python library) implementation of the Faster RCNN architecture. The agenda of the training process is to minimise a loss function which depicts the error in prediction in comparison to the ground truth. On each iteration of training, the weights/parameters in the network are adjusted to minimize loss of function.

The adjustment in these weights lead to a combination of equations which when given an image, output a set of rectangular coordinates with their objectness scores respective classes, here the micronucleus. These coordinates are then mapped on the image to show the detections during inference.

#### 3.3.1 Method of evaluating the proposed flow

The authors aim to showcase the inherent ability of the proposed model to learn and identify morphological features of MN. Instead of just mentioning results for one such instance with a predefined dataset distribution into train, validate and test sets, a n-fold validation approach was chosen. A n-fold validation study helps us gauge the performance of the framework instead of showing results for one instance. Thus, generated results should be considered to hold a more generic value showcasing the learning ability of this framework in the task of MN detection. For a 3-fold validation (n=3), the annotated dataset is divided into 3 parts randomly of equal size. Two parts are used for training purposes and the results are quoted on tests on the third part. The training part is divided into training set (90%) and the validation set (10%). Each part is tested once with the other two being used for training. The results shown here are an average of 3 cross-validation results.

#### 3.3.2 Evaluation metrics

Intersection over union (IoU) is used to measure the overlap between the predicted object boundary and the ground truth object boundary to decide if the predicted object boundary is true positive or false positive or false negative. In this experiment the IoU threshold was set to 0.5 empirically. The annotations done by the annotators where their decisions coincided were identified for IoU threshold selection. We observed that the IoU of 0.5 was sufficient to ensure that boxes drawn by different annotators denoted the same object. The same margin was hence also given to the AI models in detecting micronuclei.

**Figure 3:**
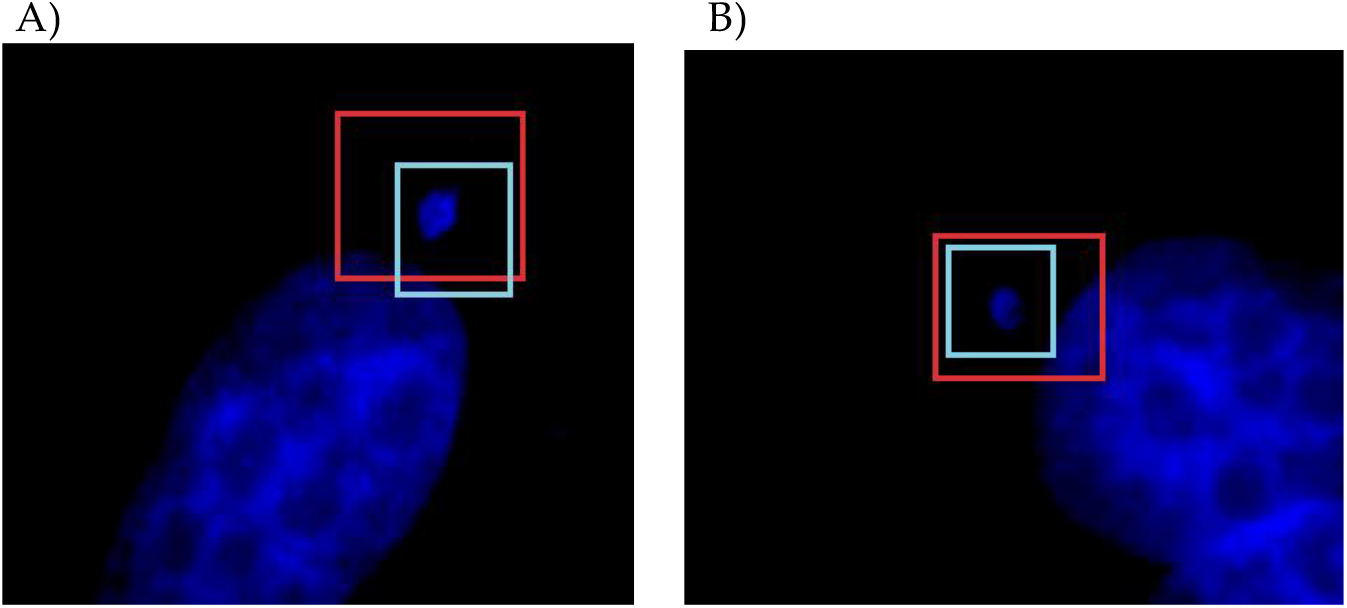
A and B are 2 different images annotated by 2 different annotators. The red box are the annotations done by annotator-1 and the blue boxes are by annotator −2. The IoU for these boxes were calculated over all the annotations. The average of those values was used to determine the IoU threshold of 0.5 used for the AI model.

Precision, recall and mean average precision (mAP) were used as metrics to gauge the results of this study. mAP is a popular metric to gauge the performance of an object detection model. It is calculated by finding the area under the precision-recall curve and averaging over all the classes.

## 4. Results

Micronuclei, the well-studied biomarker of genome instability can be induced due to dysfunctional genes and genotoxic agents. For imaging of MN, U-2OS cells were cultured with low concentrations of aphidicolin known to induce replication stress, the key driving factor of chromosome instability in dividing cells. The presence of binucleated cells represent cytokinesis block and aphidicolin treated cells displayed increased number of MN as compared to untreated cells (Fig 4). Little to no cellular debris were identified in the areas analysed under microscope. Three or more MN were excluded from scoring to avoid artefacts du1642e to catastrophic cellular events such as necrosis. To further ensure the specificity of our results, only DAPI-stained binucleated with distinct micronuclei in the vicinity were scored (Fig 5).

**Figure 4.**
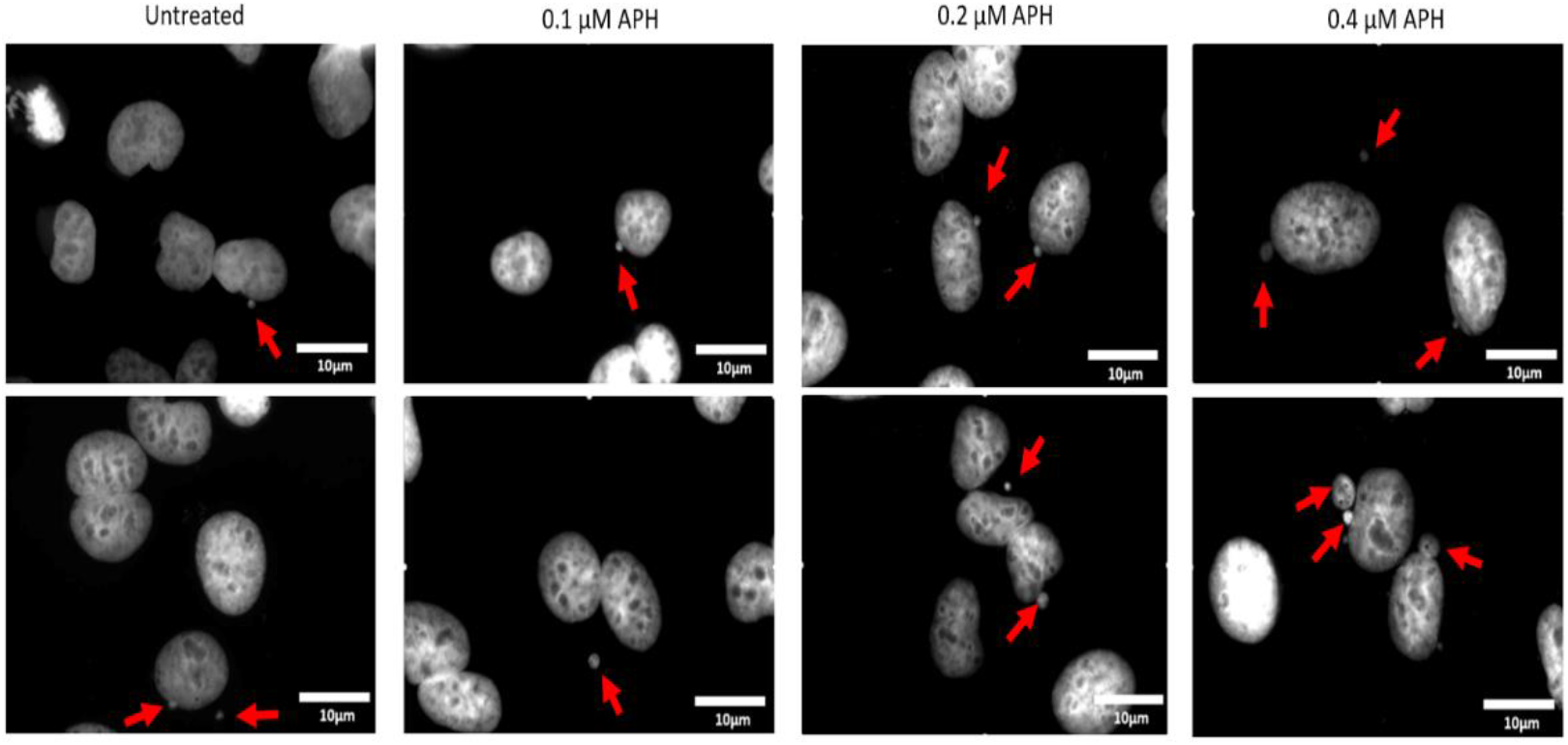
Formation of micronuclei after aphidicolin-induced replication stress. Representative images in grey scale indicate nuclei of human osteosarcoma cells cultured in the presence or absence of 0.1, 0.2 and 0.4 μM DNA polymerase inhibitor aphidicolin. Binucleated cells indicate cytokinesis block by the addition of cytochalasin B, red arrows indicate micronuclei.

**Figure 5.**
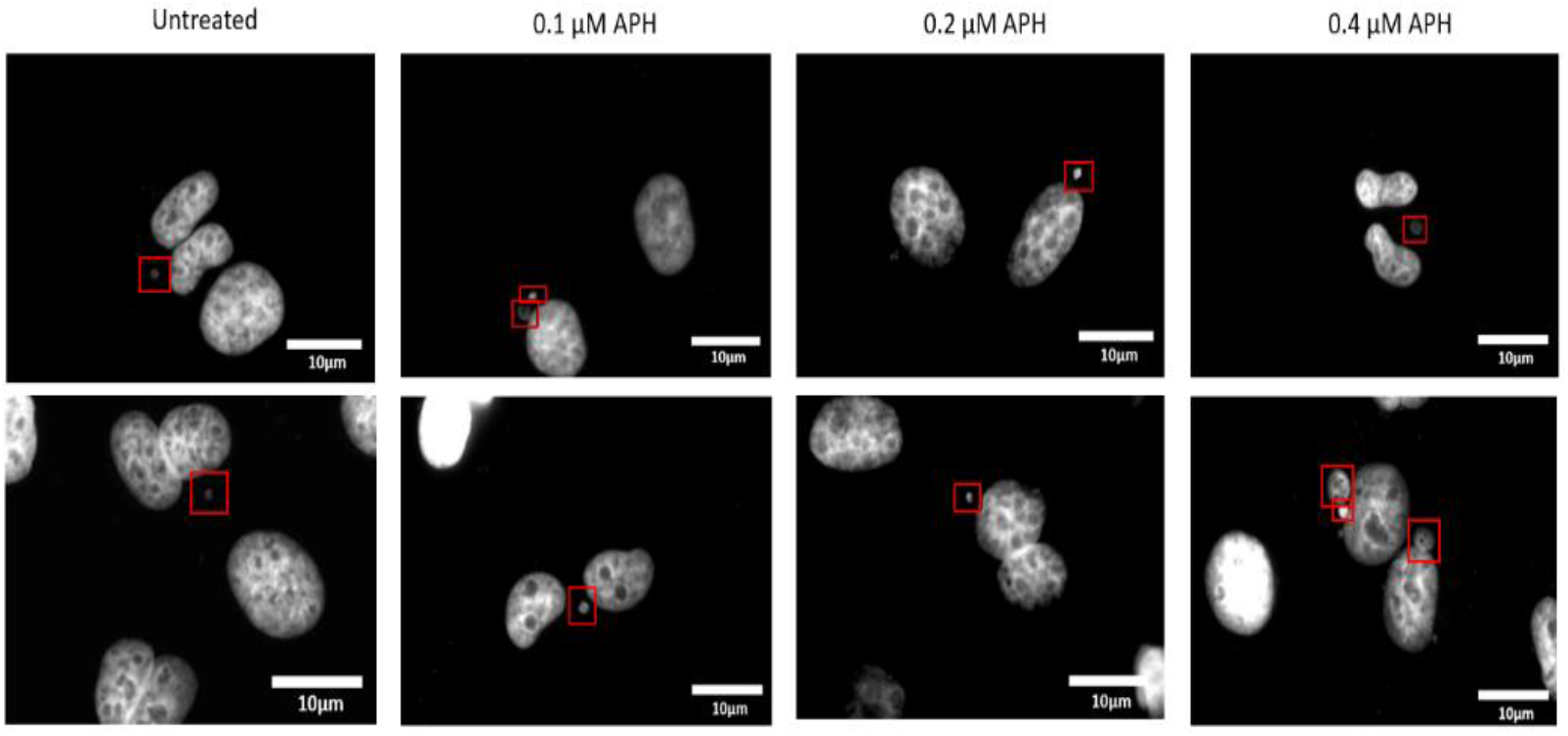
Scoring of micronuclei in a cytokinesis-block micronucleus assay. Cells were cultured on coverslips and treated with 0.1-0.4 μM aphidicolin for a period of 48 hrs before staining with DNA binding blue, fluorescent dye DAPI. Representative grey scale images indicate cells blocked at anaphase stage of cell cycle with 2 daughter nuclei of similar size in proximity to each other. Red boxes indicate, micronuclei identified by manual/ AI scoring.

Overall, the proposed workflow for deep learning framework proposed here exhibited acceptable performance in the detection of MN. This being an object detection task, mAP was considered as the primary metric. The AI model was able to reach a mAP of 0.93 at a confidence threshold (objectness score) of 0.9, a higher threshold can be correlated with a higher confidence. The results are as follows: sensitivity: 86%, precision: 78%; mAP: 0.93(93%). From figure 6, it is evident that the mAP, sensitivity, and specificity remained almost the same or increased as the threshold was increased, which signifies the reliability of the AI decisions.

**Figure 6.**
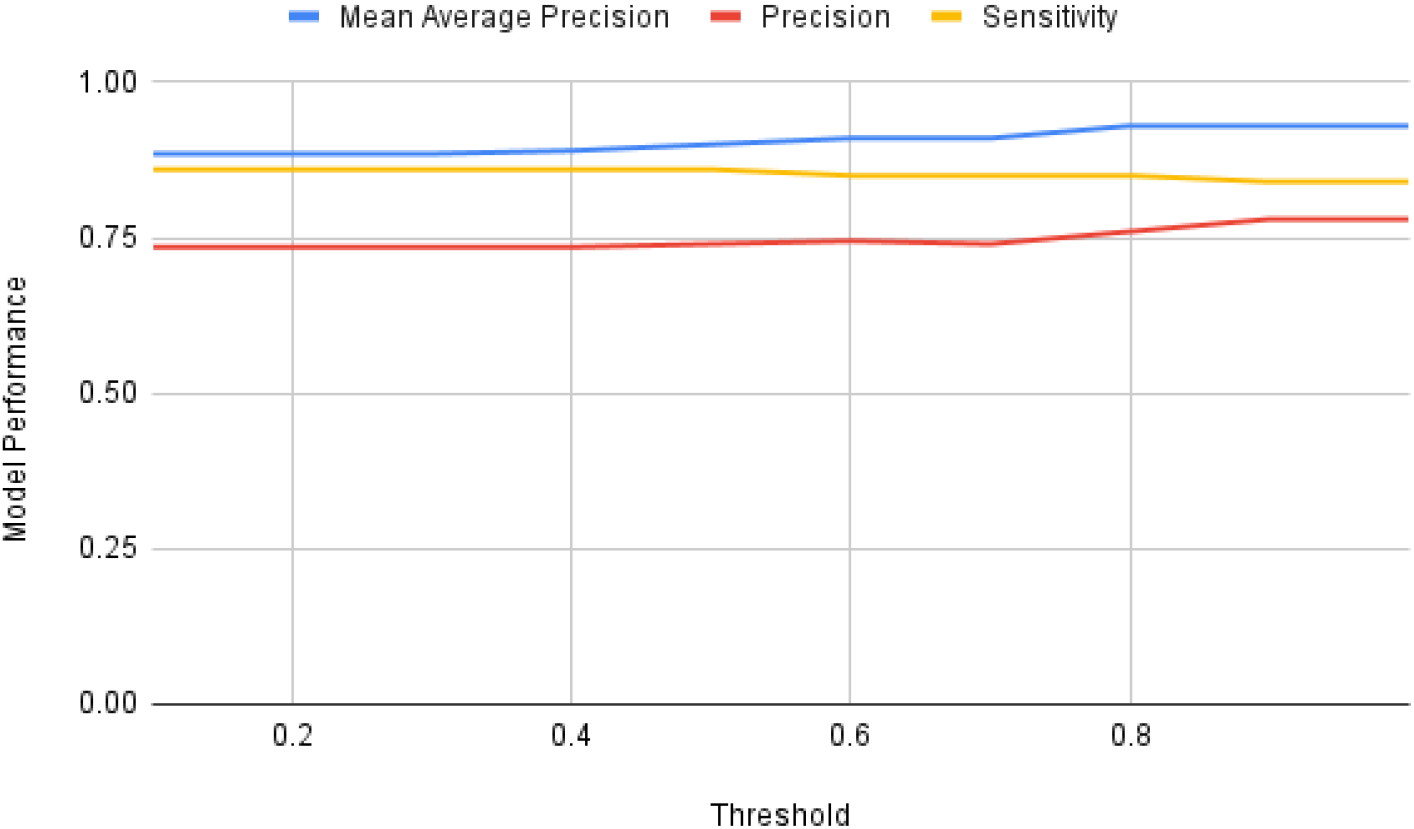
Graph depicting the model performance using metrics like mAP, precision and sensitivity (y-axis) against the confidence threshold (x-axis: objectness score) of the detection (0.1-1, on a scale of 0 being least confident to 1 being most confident).

## 5. Discussion

*In vitro* and *in vivo* cytogenetic tests are used in epidemiological studies and in the investigation of biomarkers of human diseases. The widely used micronucleus test method requires manual scoring of MN. Previous studies [19] have discussed the difficulty faced by researchers in the detection of MN in fluorescent CBMN assay images. These results were broadly driven by the difficulty of differentiating MN from debris, other artefacts, and human error.

The proposed method in this study was able to detect MN with a high efficacy. Sample preparation methodologies used in the study ensured the minimum occurrence of debris/artefacts. Further studies will be required to assess the performance of the model under scenarios of excess debris in samples. For object detection use cases, mAP is the integrated score that tells the complete story of an object detection model. The authors were not able to find another study that talked in terms of this metrics. Thus, the sensitivity and precision metrics have been included in the results to get a sense of the performance of the proposed framework. Bahreyni Toossi et al reported a sensitivity of 82% for MN detection in the detected BN [24]. The current study yielded similar sensitivity scores in a more challenging environment of detecting individual MN on the images of a slide instead of detecting the presence of MN in a detected cell image compared to previous reports [24].

The AI model was trained with relatively small data however the results presented here show potential to decrease the amount of manual work currently undertaken by researchers working in laboratories involved in delineating the link between genomic instability and molecular basis of diseases. Such methodologies when integrated will play an essential role in standardizing and speeding up the process of detection and counting of MN. Improvements have been observed in previous AI studies with enhanced data quality and size. The proposed AI model was also tested on a similar public dataset BBBC039^1^, and exhibited potential to detect the MN in these images however the accuracy cannot be commented in this study.

As the pace of sequencing genomes increases, the quantities of raw data produced by next generation sequencing methods has increased massively [18]. Over the past 4 decades many bioinformatics tools have been developed that enable the analysis and investigation of sequenced data. ML techniques could be utilised to further enhance existing bioinformatics tools and to develop entirely new classes of bioinformatics tools that leverage auto-ml platforms such as Logy.AI for investigation of genome instabillity. In theory, any repetitive activity which involves a human having to analyse images and count specific shapes - or more generally *patterns* (which could also include textual genetic sequences) - could benefit from being automated using an ML platform like Logy.AI. In fact, this may indeed become a necessity as data volumes continue to increase and human resources remain in short supply. By using no code ML training, the writing of custom application specific software is eliminated. The advantages of this include reduced costs, reduced IT infrastructure, and reduced IT training costs as non-programmer bio-scientists can train an AI’s without learning to code. AI model training by LaiAMLP was performed on an Intel®Xeon @ 2.70GHz machine (aws p2.xlarge [30]) with an NVIDIA®Tesla K80 [31] with 12 GB of memory running Ubuntu 18.04 [32]optimised for deep-learning. It took 4 hours to train each model. The total computational cost of training the model was ~10USD*^2^. The resultant model can be run on a personal computer and the running cost post the creation of the AI model is thus very minimal. This demonstrates that platforms like LaiAMLP are very cost effective in building such AI models on demand for labs of all sizes and domains.

There are several issues in manual counting and annotation of micronuclei in fluorescent CBMN assay images. There is also the additional problem of researcher *bias* in identification of artefacts leading different results. However, the bias present in a researcher can also be passed on to the ML (known as *machine bias*) [19]. Thus, it is encouraged to use more than one human during training the AI, providing a ‘consensus’ of what is positive and negative. In addition, after the AI has been trained, it is still important to randomly sample ‘counted’ images to determine accurate counting, and ‘tweak’ the AI if required, essentially providing more learning opportunity and thus more future accuracy. Regular testing of the workflow is essential to ensure the performance of the system to acceptable standards and thresholds.

## 6. Limitations and Future Scope

This study has attempted to explore a new way of outsourcing some skills to carefully crafted AI models to ease the work of researchers and increase the speed of research. There are a few perceived limitations to this study which opens doors for future research. The dataset used is still small, repeating this study with larger-sized data will help establish the efficacy of such a system and to shed light on the overall time saved and gauge the performance of a human-AI duo system in detecting micronuclei. Blur objects around the parent nuclei constituted a major portion of cases where the model faced difficulties in differentiating between micronuclei and a possible artefact. A possible solution could be to use bigger boxes to include the colouring of the neighbouring nuclei during annotations enabling the model to use colour intensity of the surrounding nuclei to rule out such artefacts. Colour intensity of MN are similar to that of the neighbouring nuclei and are not too far from the parent nuclei while artefacts/debris on the other hand exhibit a relatively different colour intensity. The authors believe that this will be a very important future scope to explore: to include the surrounding/contextual information in the conventional detection process.

## 7. Conclusion

In the current work, we demonstrate how advancements in the field of computer science, especially in AI [33,34], can be leveraged in genome instability studies to build viable tools. The presented work is fully working in terms of functional proof of concept and is intended to provide motivation to encourage further work in building such AI powered tools. The AI models proposed here were able to achieve convincing results for the detection of MN in CBMN assay images with a mean average precision of > 90%. There is a potential use of this as a suggestive tool to speed up the counting of MN which is predominantly carried out manually today.

ML techniques could be utilised to further enhance existing bioinformatics tools and to develop entirely new classes of bioinformatics tools that leverage AI as well as research in biological sciences. In theory, any repetitive activity which involves a human having to analyse images and count specific shapes- or more generally *patterns* not limited to images could benefit from being automated using AI. The sheer speed of building and deploying such cost-effective systems will play a pivotal role in revolutionising modern research. This will aid the professionals to work on more pressing matters and let the machine do the mundane tasks both more accurately and reliably.

## Author Contributions

Conceptualization, Kalpana Surendranath (K.S) and Anand Panchbhai (A.P); writing-original draft preparation, K.S., A.P., Munuse Ceyda Ishanzadeh (M.C.I);, DNA damage studies, A.S.,N.S.S.,R.K.; Testing the model and visualisation M.C.I and A.P. supervision, J.J.M. and K.S. All authors have read and agreed to the published version of the manuscript.

## Data Availability Statement

Two datasets were analysed as part of this study CBMNA2021 was specifically curated for the purpose of this study and is available from the corresponding authors on request. The second dataset used in this study was released as part of another study and can be found at https://bbbc.broadinstitute.org/BBBC039.

## Acknowledgments

We gratefully acknowledge the collaborative efforts of members of Genome Engineering laboratory in the establishment of CRISPR knockout models. We thank Khalid Akram for extending his support in critical reading of the manuscript.

## Footnotes

1 https://bbbc.broadinstitute.org/BBBC039

2 The price mentioned here only comprises the computational cost of AWS as on 11th July 2021.

## References

1. Hoeijmakers, J.H.J. DNA Damage, Aging, and Cancer. N. Engl. J. Med. 2009, 361, 1475–1485, doi:10.1056/NEJMra0804615.

2. Bonassi, S.; Coskun, E.; Ceppi, M.; Lando, C.; Bolognesi, C.; Burgaz, S.; Holland, N.; Kirsh-Volders, M.; Knasmueller, S.; Zeiger, E.; et al. The HUman MicroNucleus Project on eXfoLiated Buccal Cells (HUMN(XL)): The Role of Life-Style, Host Factors, Occupational Exposures, Health Status, and Assay Protocol. Mutat. Res. 2011, 728, 88–97, doi:10.1016/j.mrrev.2011.06.005.

3. Capper, D.; Jones, D.T.W.; Sill, M.; Hovestadt, V.; Schrimpf, D.; Sturm, D.; Koelsche, C.; Sahm, F.; Chavez, L.; Reuss, D.E.; et al. DNA Methylation-Based Classification of Central Nervous System Tumours. Nature 2018, 555, 469–474, doi:10.1038/nature26000.

4. Kwon, M.; Leibowitz, M.L.; Lee, J.-H. Small but Mighty: The Causes and Consequences of Micronucleus Rupture. Exp. Mol. Med. 2020, 52, 1777–1786, doi:10.1038/s12276-020-00529-z.

5. Samwer, M.; Schneider, M.W.G.; Hoefler, R.; Schmalhorst, P.S.; Jude, J.G.; Zuber, J.; Gerlich, D.W. DNA Cross-Bridging Shapes a Single Nucleus from a Set of Mitotic Chromosomes. Cell 2017, 170, 956–972.e23, doi:10.1016/j.cell.2017.07.038.

6. Di Marco, S.; Hasanova, Z.; Kanagaraj, R.; Chappidi, N.; Altmannova, V.; Menon, S.; Sedlackova, H.; Langhoff, J.; Surendranath, K.; Hühn, D.; et al. RECQ5 Helicase Cooperates with MUS81 Endonuclease in Processing Stalled Replication Forks at Common Fragile Sites during Mitosis. Mol. Cell 2017, 66, 658–671.e8, doi:10.1016/j.molcel.2017.05.006.

7. Delahunt, C.B.; Jaiswal, M.S.; Horning, M.P.; Janko, S.; Thompson, C.M.; Kulhare, S.; Hu, L.; Ostbye, T.; Yun, G.; Gebrehiwot, R.; et al. Fully-Automated Patient-Level Malaria Assessment on Field-Prepared Thin Blood Film Microscopy Images. 2019 IEEE Global Humanitarian Technology Conference (GHTC) 2019.

8. Website Available online: https://doi.org/10.48550/arXiv.2011.14329.

9. Lakhani, P.; Sundaram, B. Deep Learning at Chest Radiography: Automated Classification of Pulmonary Tuberculosis by Using Convolutional Neural Networks. Radiology 2017, 284, 574–582, doi:10.1148/radiol.2017162326.

10. Alzubaidi, L.; Fadhel, M.A.; Al-Shamma, O.; Zhang, J.; Duan, Y. Deep Learning Models for Classification of Red Blood Cells in Microscopy Images to Aid in Sickle Cell Anemia Diagnosis. Electronics 2020, 9, 427.

11. Karbhari, Y.; Basu, A.; Geem, Z.-W.; Han, G.-T.; Sarkar, R. Generation of Synthetic Chest X-Ray Images and Detection of COVID-19: A Deep Learning Based Approach. Diagnostics (Basel) 2021, 11, doi:10.3390/diagnostics11050895.

12. Krause, J.; Gulshan, V.; Rahimy, E.; Karth, P.; Widner, K.; Corrado, G.S.; Peng, L.; Webster, D.R. Grader Variability and the Importance of Reference Standards for Evaluating Machine Learning Models for Diabetic Retinopathy. Ophthalmology 2018, 125, 1264–1272, doi:10.1016/j.ophtha.2018.01.034.

13. Lucius, M.; De All, J.; De All, J.A.; Belvisi, M.; Radizza, L.; Lanfranconi, M.; Lorenzatti, V.; Galmarini, C.M. Deep Neural Frameworks Improve the Accuracy of General Practitioners in the Classification of Pigmented Skin Lesions. Diagnostics (Basel) 2020, 10, doi:10.3390/diagnostics10110969.

14. Jacobs, C.; van Ginneken, B. Google’s Lung Cancer AI: A Promising Tool That Needs Further Validation. Nature Reviews Clinical Oncology 2019, 16, 532–533.

15. Tschandl, P.; Rinner, C.; Apalla, Z.; Argenziano, G.; Codella, N.; Halpern, A.; Janda, M.; Lallas, A.; Longo, C.; Malvehy, J.; et al. Human-Computer Collaboration for Skin Cancer Recognition. Nat. Med. 2020, 26, 1229–1234, doi:10.1038/s41591-020-0942-0.

16. Dias, R.; Torkamani, A. Artificial Intelligence in Clinical and Genomic Diagnostics. Genome Medicine 2019, 11.

17. Rodrigues, M.A.; Probst, C.E.; Zayats, A.; Davidson, B.; Riedel, M.; Li, Y.; Venkatachalam, V. The in Vitro Micronucleus Assay Using Imaging Flow Cytometry and Deep Learning. NPJ Syst Biol Appl 2021, 7, 20, doi:10.1038/s41540-021-00179-5.

18. Alafif, T.; Qari, S.; Albassam, A.; Alrefaei, A. Deep Transfer Learning for Nucleus and Micronucleus Recognition. 2020 First International Conference of Smart Systems and Emerging Technologies (SMARTTECH) 2020.

19. Caicedo, J.C.; Roth, J.; Goodman, A.; Becker, T.; Karhohs, K.W.; Broisin, M.; Molnar, C.; McQuin, C.; Singh, S.; Theis, F.J.; et al. Evaluation of Deep Learning Strategies for Nucleus Segmentation in Fluorescence Images. Cytometry A 2019, 95, 952–965, doi:10.1002/cyto.a.23863.

20. Carpenter, A.E.; Jones, T.R.; Lamprecht, M.R.; Clarke, C.; Kang, I.H.; Friman, O.; Guertin, D.A.; Chang, J.H.; Lindquist, R.A.; Moffat, J.; et al. CellProfiler: Image Analysis Software for Identifying and Quantifying Cell Phenotypes. Genome Biol. 2006, 7, R100, doi:10.1186/gb-2006-7-10-r100.

21. Reuter, J.A.; Spacek, D.V.; Snyder, M.P. High-Throughput Sequencing Technologies. Molecular Cell 2015, 58, 586–597.

22. Tarca, A.L.; Carey, V.J.; Chen, X.-W.; Romero, R.; Drăghici, S. Machine Learning and Its Applications to Biology. PLoS Comput. Biol. 2007, 3, e116, doi:10.1371/journal.pcbi.0030116.

23. Su, H.-H.; Pan, H.-W.; Lu, C.-P.; Chuang, J.-J.; Yang, T. Automatic Detection Method for Cancer Cell Nucleus Image Based on Deep-Learning Analysis and Color Layer Signature Analysis Algorithm. Sensors 2020, 20, 4409, doi:10.3390/s20164409.

24. Bahreyni Toossi, M.T.; Azimian, H.; Sarrafzadeh, O.; Mohebbi, S.; Soleymanifard, S. Automatic Detection of Micronuclei by Cell Microscopic Image Processing. Mutat. Res. 2017, 806, 9–18, doi:10.1016/j.mrfmmm.2017.07.012.

25. Shen, X.; Chen, Y.; Li, C.; Yang, F.; Wen, Z.; Zheng, J.; Zhou, Z. Rapid and Automatic Detection of Micronuclei in Binucleated Lymphocytes Image. Sci. Rep. 2022, 12, 1–14, doi:10.1038/s41598-022-07936-4.

26. Ren, S.; He, K.; Girshick, R.; Sun, J. Faster R-CNN: Towards Real-Time Object Detection with Region Proposal Networks. 2015, doi:10.48550/arXiv.1506.01497.

27. Huang, J.; Rathod, V.; Sun, C.; Zhu, M.; Korattikara, A.; Fathi, A.; Fischer, I.; Wojna, Z.; Song, Y.; Guadarrama, S.; et al. Speed/accuracy Trade-Offs for Modern Convolutional Object Detectors. 2016, doi:10.48550/arXiv.1611.10012.

28. He, K.; Zhang, X.; Ren, S.; Sun, J. Deep Residual Learning for Image Recognition. 2015, doi:10.48550/arXiv.1512.03385.

29. Russakovsky, O.; Deng, J.; Su, H.; Krause, J.; Satheesh, S.; Ma, S.; Huang, Z.; Karpathy, A.; Khosla, A.; Bernstein, M.; et al. ImageNet Large Scale Visual Recognition Challenge. Int. J. Comput. Vis. 2015, 115, 211–252, doi:10.1007/s11263-015-0816-y.

30. Website available online: https://docs.aws.amazon.com/AWSEC2/latest/APIReference/Welcome.html.

31. Nickolls, J.; Buck, I.; Garland, M.; Skadron, K. Scalable Parallel Programming with CUDA. Queue 2008, 6, 40–53.

32. Sobell, M.G. A Practical Guide to Linux® Commands, Editors, and Shell Programming; Pearson Education India, 2006; ISBN 9788131726501.

33. Ting, D.S.W.; Liu, Y.; Burlina, P.; Xu, X.; Bressler, N.M.; Wong, T.Y. AI for Medical Imaging Goes Deep. Nat. Med. 2018, 24, 539–540, doi:10.1038/s41591-018-0029-3.

34. Kundu, S.; Elhalawani, H.; Gichoya, J.W.; Kahn, C.E. How Might AI and Chest Imaging Help Unravel COVID-19’s Mysteries? Radiology: Artificial Intelligence 2020, 2, e200053.

